# Kia kaua te reo e rite ki te moa, ka ngaro: Do not let the language suffer the same fate as the moa

**DOI:** 10.1101/817148

**Authors:** Tessa Barrett-Walker, Michael J. Plank, Rachael Ka’ai-Mahuta, Daniel Hikuroa, Alex James

## Abstract

More than a third of the world’s languages are currently classified as endangered and more than half are expected to go extinct by 2100. Strategies aimed at revitalising endangered languages have been implemented in numerous countries, with varying degrees of success. Here, we develop a new model regarding language transmission by dividing the population into defined proficiency categories and dynamically quantifying transition rates between categories. The model can predict changes in proficiency levels over time and, ultimately, whether a given endangered language is on a long-term trajectory towards extinction or recovery. We calibrate the model using data from Wales and show that the model predicts that the Welsh language will thrive in the long term. We then apply the model to te reo Māori, the Indigenous language of New Zealand, as a case study. Initial conditions for this model are estimated using New Zealand census data. We modify the model to describe a country, such as New Zealand, where the endangered language is associated with a particular subpopulation representing the Indigenous People. We conclude that, with current learning rates, te reo Māori is on a pathway towards extinction, but identify strategies that could help restore it to an upward trajectory.

## Introduction

The world has an estimated 7000 languages, of which over 2800 are endangered (1). Between 50% and 90% of the world’s languages are expected to be extinct by 2100 (2). Many languages have become endangered as a result of colonisation, associated with oppression of Indigenous Peoples and cultural subjugation, in a pattern repeated around the world since the early 17^th^ century, including Indigenous languages in North and South America, Africa, and Australia (1, 3, 4). Examples of this pattern include Hawai’i and Aoteroa (New Zealand), where the Indigenous languages have followed almost identical patterns of decline (5–7). In New Zealand, the number of speakers of the Indigenous language, te reo Māori, was systematically reduced in the colonial period. This was done via numerous means including prohibition of te reo Māori in schools, stigmatisation of Māori culture, severance of intergenerational transmission, the Tohunga Suppression Act of 1907, and government relocation policy (8–10). Te reo Māori is classified as Threatened according to the Expanded Graded Intergenerational Disruption Scale (11). Te reo is intrinsically linked to all that Māori culture encompasses (12, 13) and suffered greatly to the verge of extinction in the 1960s and 70s (9, 12). There is a widespread sense that to lose one’s language is to lose one’s culture and identity (10, 14, 15).

More recently, some countries have begun to recognise the effects of their history of cultural oppression on language endangerment. Language revitalisation strategies have been implemented, with varying degrees of success. These include such measures as: affording official language status (8); integration into school curricula (16); bilingual or language-immersion schooling (10, 16–18); mentor-apprentice programs (19); support for language media (20); strategies to re-establish and maintain intergenerational transmission (21, 22); and strategies to promote engagement with an endangered language (18). In 2018, the New Zealand Government set a target that, by 2040, one million New Zealanders (approximately 20% of the population) will be able to speak at least basic te reo Māori, and it will be the primary language for 150,000 Māori (18). The ultimate success or failure of such strategies in revitalising an endangered language will depend on their effectiveness in reversing the centuries-long trend of decline in the number of users of that language.

Most work on language change is done by historical or socio-linguistics (17, 23–26). This body of work has provided the theoretical and empirical foundation for mathematical modelling approaches, which broadly fall into two types. The first is concerned with the evolution of language itself, for example changes in a language’s phonetic, semantic or syntactic features over time (27–30). The process of language change has also been studied in the context of cultural evolution (31–33). The second type of modelling approach, into which this study falls, assumes that the language is fixed and models trends in the number of speakers of that language over time due to shifts in individuals’ language use. This approach was popularised by Abrams and Strogatz (34), who assumed that two languages compete for speakers and an increase in the number of speakers or in the perceived status of one language increases its attractiveness. In this model, one language will ultimately dominate and push the other to extinction. It was subsequently shown that including other factors, such as spatial heterogeneity modelled by a reaction-diffusion equation or population dynamics modelled by a reproduction term, could allow the languages to coexist (35–37). Spatially explicit models allow the spread or regression of a language over time to be investigated, and have been applied to the geographic range of Gaelic in Scotland (38, 39) and Slovenian in Austria (40). Other extensions to (34) consider a wider range of societal conditions and parameters including bilingualism and intergenerational transmission (30, 41–43). Modelling based on these works has been applied to te reo Māori by (44), where the amount of te reo Māori heard, family contribution and community contribution were recognised as influential factors. In addition to these differential equation models, agent-based models have been used to add finer-scale information and environmental factors (40) onto the same underlying mechanism of competition between a dominant and a minority language. Mathematically, these models are of Lotka-Volterra type with a mixture of competitive and predator-prey interactions (43, 45).

It is certainly true that language extinction is typically caused by loss of speakers to other languages rather than extinction of the population of speakers (10, 39, 46). One-way bilingualism, whereby speakers of an endangered language typically also speak the majority language but not vice versa, can also be a threat for a minority language (47, 48). Nevertheless, most endangered languages will only be able to survive and flourish by coexisting with other languages via bilingual or multilingual speakers and multiple language domains (24, 49–51), rather than outcompeting them (3). This requires that languages are accorded equal socioeconomic status (23, 42, 48).

With any language, there are varying degrees of proficiency of language use. The Common European Framework of Reference for Languages (CEFR) (52) categorises language proficiency into three broad levels: basic, independent and proficient. Basic users are able to communicate, with assistance and in simple terms, matters related to their immediate self, actions and environment. Independent users can understand and communicate experiences, explanations and opinion. Proficient users can communicate fluently, express themselves in all circumstances and understand complex or implicit meanings within text and conversation (52).

In this study, we explore factors affecting the survival of an endangered language by developing a dynamic model which divides the population by level of proficiency with the language. The model has three important differences from those described above. (I) It does not treat the endangered language as being in direct competition with the dominant language, but instead assumes that the vast majority of individuals are competent in the dominant language, regardless of whether they learn the endangered language. (II) It considers differences in proficiency level with the endangered language across the population. (III) It explicitly includes two language acquisition routes, namely intergenerational transmission and language learning in school-age children or adults. Therefore, state changes in the model do not correspond to transitions of individuals between competing languages (as in (34, 35)) and/or a bilingual state (42, 43), but to transitions between different levels of proficiency with the endangered language, independently from the dominant language. The proficiency levels in the model are based loosely on the CEFR categories. The exact choice of scale used to define proficiency categories is not important and similar models could be built based on alternative scales. Our approach has some similarities with the ZePA model of language revitalisation, which classifies people into either a zero, passive or active state(46, 53). In addition to language proficiency, these states encompass people’s values, attitudes and behaviour. We take a simplified approach of defining the model categories solely on an individual’s level of language proficiency. However, unlike the ZePA model, we attempt to quantify transitions between categories as a result of language acquisition and basic demographic processes. We do this via a system of differential equations that describe changes in the numbers of people in each category. The model is a coarse-scale model for processes affecting the number of people with varying levels of language proficiency at a population level. The model neglects variations among individuals in the myriad of factors that can affect learning outcomes, for example: access to learning resources or other language media (20); learning biases (33, 54); geographical variations (40); domains of language use (49, 51); attitudes towards the language itself (53). Nor does it attempt to account for historical processes and events. Instead, the aim is to predict whether the language is on trajectory towards extinction or recovery, and to explore how different national interventions may affect this trajectory.

We parameterise the model using data from the recent resurgence of the Welsh language. Significant development in bilingual and Welsh-medium education and the presence of the language throughout the public and private sectors have positively contributed to an increase in the number of Welsh speakers (55). The Welsh language use in Wales surveys (55–57) contain categorical proficiency data and the three available surveys provide a useful set of longitudinal data over an extended time period. The context for the model in te reo Māori in New Zealand is different from that in Wales, partly because New Zealand contains relatively well defined Māori and non-Māori subpopulations. We recalibrate the model for te reo Māori using the limited volume of census data available. We use the model to assess the current trajectory of the language and to estimate the rates of learning that would be needed, relative to the situation in Wales, to achieve language revitalisation. Informed by the model, we evaluate the likelihood of meeting government targets by 2040, and compare the potential strategies for promoting revitalisation.

### Model development

We divide the population into three categories according to their level of language proficiency: basic, independent and proficient. In reality, language proficiency does not fall neatly into discrete categories, but is a continuous and multifaceted spectrum. However, in most cases, data on language proficiency are not sufficient to calibrate anything other than a very simple model. Although the CEFR definition of the basic category is those with some level of capability with language, we include in this category those with no capability at all. We consider a system of differential equations for the proportion of the population in the basic *B*(*t*), independent *I*(*t*), and proficient *P*(*t*) categories, where *B*(*t*) + *I*(*t*) + *P*(*t*) = 1. We assume that the total population size is fixed by assigning each category identical per capita birth and death rates *r*. Individuals transition between categories as a result of language acquisition. We describe the model for this process below (see below).

We assume that once a proficiency level is reached it cannot be lost, i.e. language skills do not deteriorate with time. We are confident this assumption holds for those with te reo Māori as their first language and, given the drivers for te reo Māori acquisition as a second language, it is reasonable for that group as well. Although there might be some minor proficiency loss, this would not result in movement between proficiency categories.

#### Intergenerational transmission

We assume that children of parents in the basic and independent categories are born into the basic category. A fraction *α* of the children of proficient parents are born into the proficient category, the remaining fraction 1 − *α* are born into the basic category. This assumption is an approximate model for intergenerational language transmission that assumes these children acquire language proficiency from birth. This simplifies the model because it avoids the need to explicitly model a parent-infant learning route separately from other learning routes. The parameter *α* is the intergenerational transmission rate, which is a quantity that is commonly estimated in studies of endangered languages (e.g. (55, 58)).

#### Language acquisition

Individuals can progress from one proficiency level to the next by acquiring language competency. This usually depends on direct interaction with proficient language users, either as formal language teachers or as informal social contacts. It is also possible that interaction of basic users with independent users could facilitate learning or promote the idea of language acquisition, and hence accelerate the progression from the basic to the independent category. However, for simplicity we neglect this and assume that individuals in either the basic or independent category progress to the next category at a per capita rate that is proportional to the size of the proficient category, i.e. the number of people capable of acting as teachers. This results in the system of differential equations:

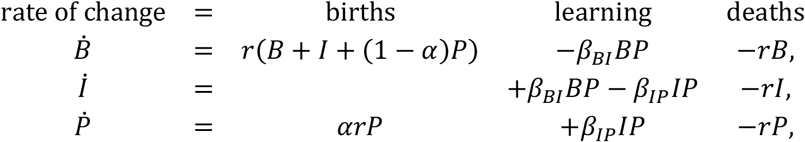

where *β*_*BI*_ and *β*_*IP*_ are learning rate parameters representing the rate of transition from the basic to the independent and from the independent to the proficient category respectively. These parameters will be affected by a number of important variables, including the quality of teachers, the motivation and commitment of learners, access to educational facilities and programmes, and the relative roles of different formal and informal learning routes. We do not attempt to account for these factors individually, but instead take a coarse-scale approach of exploring the effect of variations in the learning rate parameters *β*_*BI*_ and *β*_*IP*_.

As the total population size is assumed to be fixed, the system of equations above can be reduced to two equations for *I* and *P*:

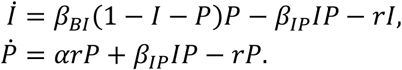

Depending on parameter values, this dynamical system has either one or three steady states. The zero steady state (*I*, *P*) = (0,0) is stable for all parameter values. When the system is in this steady state, all individuals are in the basic category and we interpret this as extinction of the language. The non-zero steady states (*I*\*, *P*\*) are given by *I*\* = (1 − *α*)*r*/*β*_*IP*_ and *P*\* is the solution of the quadratic equations

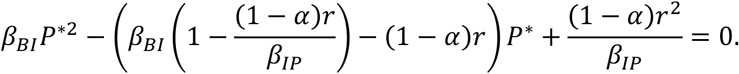

Hence, the system has a saddle-node bifurcation: below the bifurcation the only steady state at (0,0) and the language is guaranteed to go extinct from any initial condition; above the bifurcation there is an additional stable steady state with *I*\*, *P*\* > 0 and the eventual fate of the language depends on the initial conditions. Figure 2A shows the phase plane for the model in the case where the positive steady state exists. The stable manifold of the saddle steady state (red line in Fig. 2A) divides the phase space into two basins of attraction: initial conditions below the red line will result in extinction of the language; initial conditions above the red line will result in long-term survival of the language. Figure 2B shows the saddle-node bifurcation dividing the (*β*_*BI*_, *β*_*IP*_) parameter space into: (i) a region where the language will go extinct regardless of initial conditions and; (ii) a region where survival of the language is possible depending on initial conditions.

**Figure 1:**
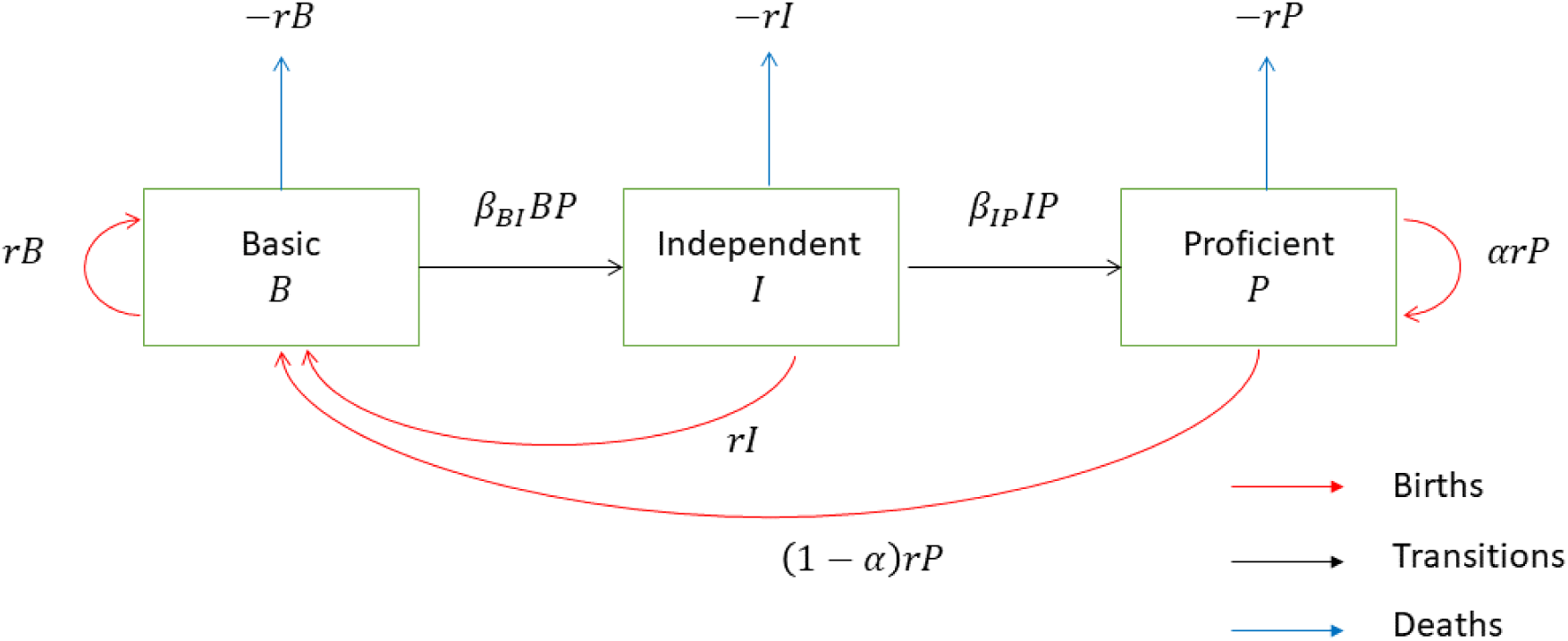
Transitions between categories occurring as a result of births, deaths and learning from proficient users.

**Figure 2.**
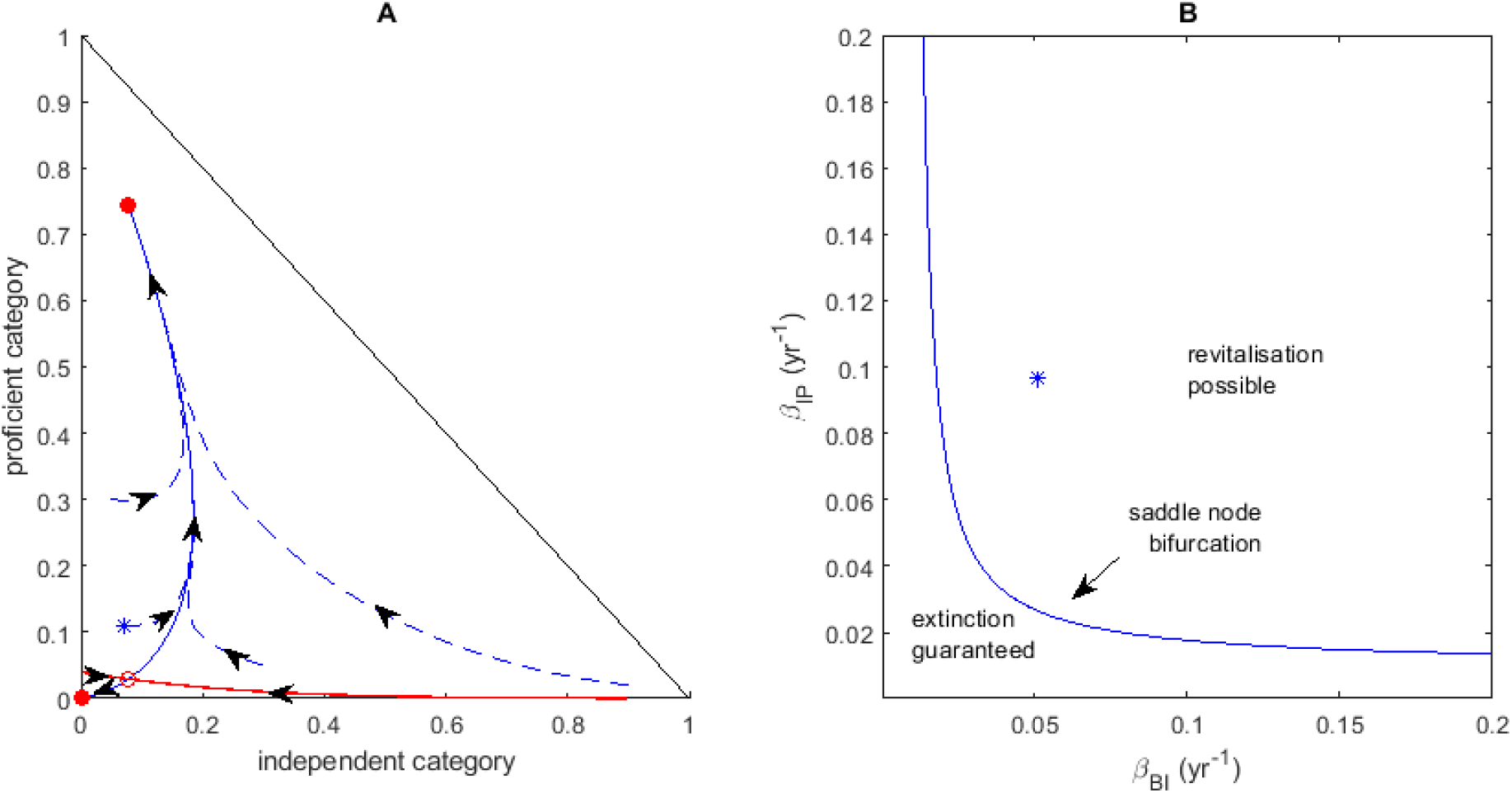
Structure of the dynamical system for the proportion of the population in the independent category (*I*) and the proficient category (*P*). (A) Trajectories (dashed blue curves) and steady states (red circles) in the (*I*, *P*) phase plane for learning rate parameters *β*_*BI*_= 0.051 yr^−1^ and *β*_*IP*_ = 0.0969 yr^−1^. For these values of *β*_*BI*_ and *β*_*IP*_, there are two positive steady states as well as the steady state at (*I*, *P*) = (0,0). The steady states at (*I*, *P*) = (0.08,0.74) and (*I*, *P*) = (0,0) are locally stable (filled red circles) and the steady state at (*I*, *P*) = (0.08,0.03) is a locally unstable saddle node (open red circle). The stable manifold of the saddle node (red curve) separates the phase space into two basins of attraction: initial conditions below the red curve tend towards the zero steady state (language extinction); initial conditions above the bold curve tend to the positive stable steady state (language revitalisation). The blue star shows the 1991 initial conditions and predicted trajectory for Welsh. (B) Two-parameter bifurcation diagram showing behaviour of the model in (*β*_*BI*_, *β*_*IP*_) parameter space. There is a saddle-node bifurcation (solid curve) separating parameter combinations for which extinction is guaranteed from those for which language revitalisation is possible. Blue star shows estimated learning rates for Welsh. Other parameter values: *α* = 0.47 and *r* = 1/70 yr^−1^.

### Model calibration and predictions for Welsh

The model has three language-related parameters: the intergenerational transmission rate *α* and the two learning rate parameters *β*_*BI*_ and *β*_*IP*_. The birth/death rate parameter *r* is purely demographic and not language-related; we set this as *r* = 1/70 yr^−1^, which corresponds to an average life expectancy of 70 years.

We use data on Welsh language users to estimate the parameters *α*, *β*_*BI*_ and *β*_*IP*_ for that context. The 1991 Welsh census found that 18.7% of the population were able to speak Welsh (59) and the 1992 Welsh Language Survey found that 61% of these individuals were fluent. Interpreting the fluent speakers as being in the proficient category, the non-fluent speakers as being in the independent category, and the non-speakers as being in the basic category, this corresponds to a system state of (*B*, *I P*) = (0.81, 0.07, 0.11) in 1991. The 2004-06 Welsh language use survey found that 20.5% of the population aged three and over were able to speak Welsh and 12% were able to speak Welsh fluently (56). This corresponds to a system state of (*B*, *I P*) = (0.795, 0.085, 0.12) in 2005. Finally the Welsh Government’s report on Welsh language use in Wales (55) categorised 24% of the population as Welsh speakers and 11% as fluent, giving (*B*, *I P*) = (0.76, 0.13, 0.11) in 2014.

The intergenerational transmission rate has been estimated for several languages including Welsh. The report on Welsh language use in Wales 2013-15 (55) found that 45% of children aged 3 to 4 could speak Welsh in households with one Welsh-speaking parent, and 82% of children aged 3 to 4 could speak Welsh in households with two Welsh-speaking parents. Assuming homogeneous mixing between different categories and 11% of the population in the proficient category, this corresponds to an average probability of intergenerational transmission of *α* = 0.47.

The learning rate parameters *β*_*BI*_ and *β*_*IP*_ represent the per capita rate at which individuals in one proficiency category progress to the next category when taught by a proficient language user. Individuals can progress from the basic to the independent category in a variety of ways, ranging from mainstream (i.e. non-immersion) schooling, social interactions with language speakers, hearing the language on the radio and television, supermarket and road signs, and enrolment in evening classes. The transition from the independent to the proficient category is more likely come about through tertiary level study, immersion schooling or family environment. Even with available data, estimating these parameters directly is difficult, so we estimate them by fitting the model to the data on (*B*, *I*, *P*) at the three time points 1991, 2005 and 2014.

We use the 1991 data to define the initial condition (see Table 1) and we then use the 2005 and 2014 data to fit the model using least squares to give *β*_*BI*_ = 0.0510 yr^−1^, *β*_*IP*_ = 0.0969 yr^−1^ (Figure 3). These fitted parameter values have a real-world interpretation: if almost all of the population were proficient, then 5.1% of the basic category would progress to independent per year, and 9.7% of the independent category would progress to proficient per year; when less than 100% of the population is proficient, these transition rates are reduced pro rata.

**Table 1.**
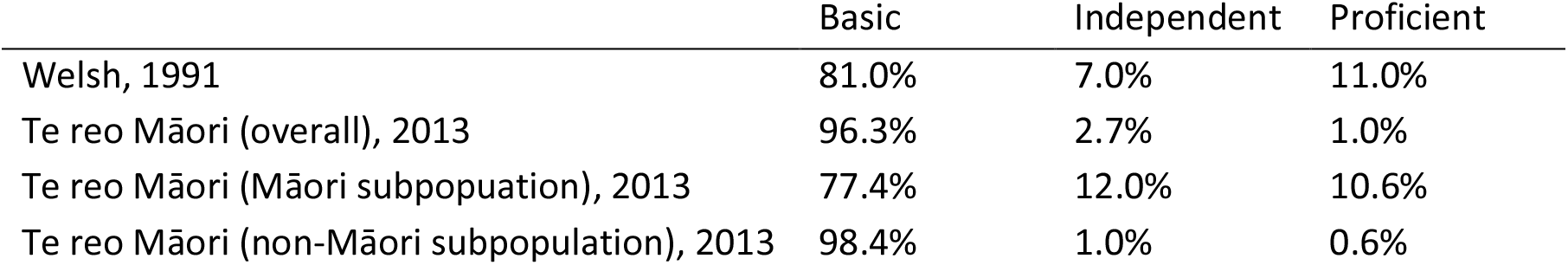
Initial conditions for the model applied to various populations, specifying the proportion of the population in the three proficiency categories.

**Figure 3:**
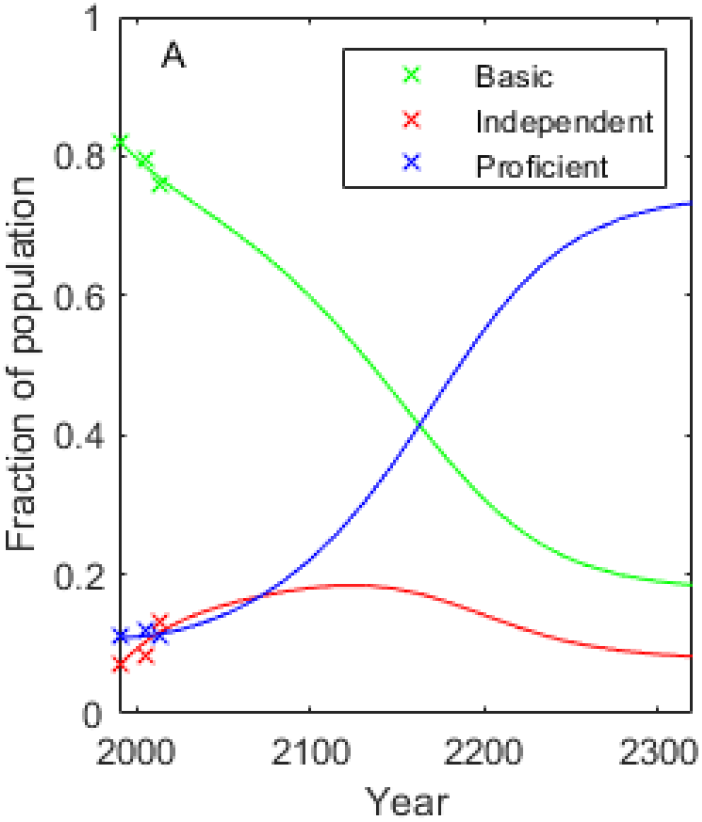
The model predicts long-term success for Welsh language revitalisation. The predicted model time series fitted to data for the Welsh language, with initial conditions taken from the first data point: *B* = 0.81, *I* = 0.07, *P* = 0.11 in 1991. Parameter values: *r* = 1/70 yr^−1^, *α* = 0.47, *β*_*BI*_ = 0.051 yr^−1^, *β*_*IP*_ = 0.0969 yr^−1^.

The model predicts that the revitalisation efforts will be successful and, in the long term, Wales will have a majority of proficient Welsh language users. Figure 2A shows that, at the estimated learning rates, provided there are at least 6% proficient speakers, almost any initial condition (including the state of the language in 1991 indicated by the blue star in Fig. 2A) would lead to successful revitalisation. The exact percentage of long-term proficient users (74% after approximately 300 years) should not be treated as a quantitatively accurate prediction. The model has been calibrated with data in a state where the majority of the population are in the basic category, and is likely to lose accuracy in very different situations as other, possibly unanticipated, factors take effect. Important among these is that the model treats the population as homogeneous, meaning that everyone has the same propensity and ability to learn, which in reality is not the case. The important model prediction is the qualitative outcome that the language is on a trajectory towards recovery. However, despite the strong long-term trend, the initial revitalisation period for the first 50 to 100 years is relatively fragile, with continued minority status and slow rates of increase, and therefore potentially sensitive to changes in learning rates or intergenerational transmission.

### Model calibration for te reo Māori

We now consider how the model can be recalibrated for te reo Māori. In countries such as New Zealand, the endangered language is associated with a relatively well-defined group of Indigenous People and the dominant language is typically associated with colonisation. It is therefore important to account for different proficiency levels in the Indigenous and non-Indigenous groups, rather than treating the population as homogeneously mixed. Within New Zealand, Māori represent approximately 15% of the total population (60), with much higher rates of reo Māori use than in the population overall. In this section, we use census data (61, 62) to estimate model parameter values and initial conditions for the population as a whole and for the Māori and non-Māori subpopulations.

Data from the 2013 survey on Māori wellbeing (62) show that, in 2013, 5% of Māori were able to speak te reo very well, 5.6% were able to speak well, 12% were able to speak fairly well, 32.1% were not able to speak very well and 45.3% were able to speak no more than a few words or phrases. We interpret the 10.6% of those able to speak very well or well as the proficient category, the 12% of those able to speak fairly well as the independent category, and the remaining 77.4% as the basic category. Of the population as a whole, 3.7% were able to speak te reo Māori (61). We estimate that 1% are in the proficient category, 2.7% in the independent category, and the remaining 96.3% in the basic category. For the non-Māori subpopulation, it is estimated that only 0.6% can speak te reo Māori (61). We err on the side of generosity and assume these are all in the proficient category and that an additional 1% of the non-Māori subpopulation are in the independent category. Table 1 shows the model initial conditions for the overall population and the Māori and non-Māori subpopulations.

(58) inferred from the 2013 census data that the intergenerational transmission rate for te reo Māori is 43.6%, close to the Welsh rate. This result encompasses Māori and non-Māori speakers of te reo.

Data for estimating the learning rate parameters from changes in proficiency levels over time are not readily available for te reo Māori. We therefore derive estimates for these parameters from the estimated Welsh values by comparing rates of participation in language learning in English-medium schools and in language-immersion schools across the two countries. Although progression to the next level of language proficiency may also occur by other means, comparing rates of participation at school-level can be used to gain an approximate estimate for learning rates in one country relative to those in another country. Although proficiency levels may differ across countries, second language learning in English-medium schools can, broadly speaking, be regarded as progression from the basic to the independent category; immersion or language-medium education can be regarded as progression from the independent to the proficient category. New Zealand’s linguistic milieu includes Polynesian languages that are grammatically and lexically similar to te reo Māori. It is possible that, in local areas with significant Pasifika populations, te reo learners experience a beneficial effect from being exposed to another Polynesian language. However, at a national scale this effect is likely to be small as these languages are used uncommonly and in restricted language domains.

Table 2 shows language learning participation rates in English-medium and language immersion schools in Wales and in New Zealand. Participation rates in New Zealand are substantially lower than in Wales, suggesting that the learning rate parameters for te reo Māori are lower than for Welsh. Table 2 shows estimates for the learning rate parameters for te reo Māori, derived by reducing the Welsh estimates pro rata by the relevant school participation rate.

**Table 2.** School participation rates and estimated learning rates for Welsh and te reo Māori. Columns 1 and 2 shows rates of participation in language learning in English-medium school and in language-immersion schools. Columns 3 and 4 show estimated learning rate parameters for the model. Estimated learning rates for Wales are obtain by fitting the model to data of changes in proficiency levels over time; estimated learning rates for New Zealand are derived from the Welsh estimates using the relative school participation rates. References for participation rate data: New Zealand (66); Wales – (16).

|  | Language learning<br>in English-medium<br>school | Language<br>immersion school | Estimated learning<br>rates (yr<sup>-1</sup>) |  |
| --- | --- | --- | --- | --- |
| | | | $\beta_{BI}$ | $\beta_{IP}$ |
| Welsh | 78% | 22% | 0.0510 | 0.0969 |
| Te reo Māori (Māori) | 31.3% | 9.6% | 0.0205 | 0.0423 |
| Te reo Māori (overall) | 21.1% | 2.5% | 0.0138 | 0.0016 |

### Model predictions for te reo Māori

We first apply the model to New Zealand’s population as a whole, and then to the Māori and non-Māori populations as two distinct, closed subpopulations. We assume that these three groups have the same intergenerational transmission rate *α*, but differ in their learning rates and initial conditions (see Table 1). To explore the potential effect of language revitalisation strategies, we present model results across a range of learning rates, using the values estimated in Table 2 as benchmarks.

Figure 4 shows the long-term, steady-state proficient population for the overall New Zealand population and the Māori and non-Māori subpopulations, for a range of learning rate parameters. The Welsh learning rates are marked with a green star; the learning rates estimated for the New Zealand population are marked with yellow stars.

**Figure 4:**
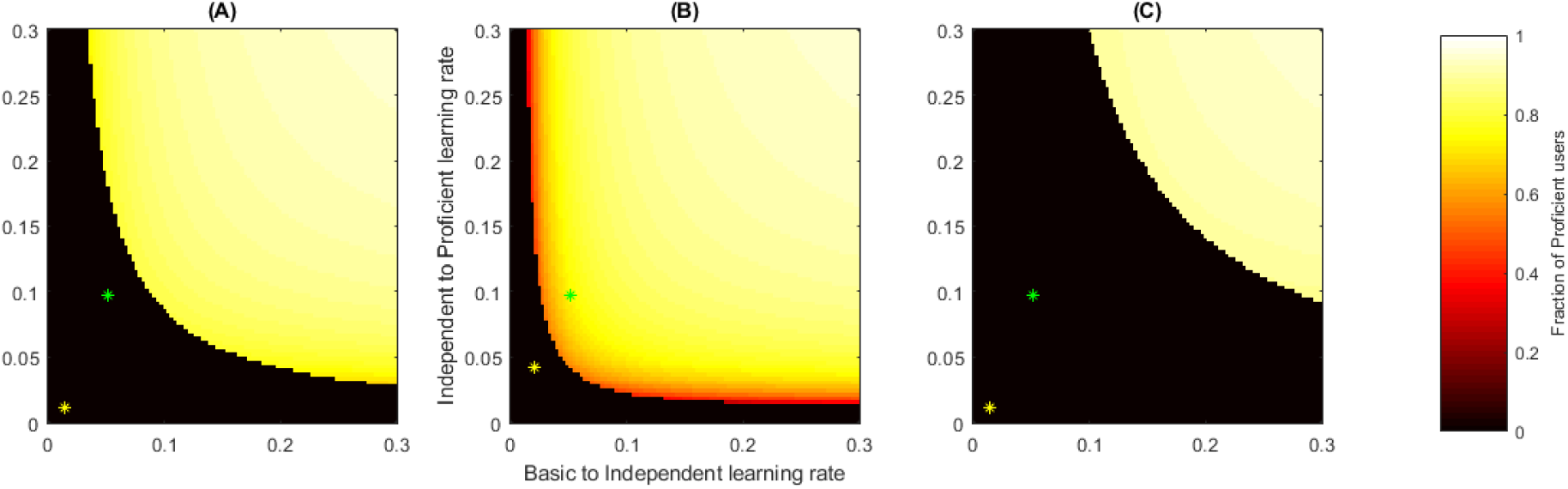
Successful revitalisation of te reo Māori is possible if the learning rate parameters are high enough. Long-term language outcomes for: (A) the New Zealand population as a whole; (B) the Māori subpopulation as a separate group; (B) the non-Māori subpopulation as a separate group. Colours show the long-term proportion of the population in the proficient category starting from initial conditions estimated for the relevant population (Table 1) and for a range of learning rate parameters *β*_*BI*_ and *β*_*IP*_. Black regions correspond to learning rate parameters that lead to long-term extinction of the language; coloured regions correspond to long-term success of the language. Green stars indicate the estimated learning rate parameters for Welsh; yellow stars indicate the estimate learning rate parameters for te reo Māori in the relevant group.

The model predicts that, when the New Zealand population is considered as a whole (Figure 4A), the language is on a path towards extinction at current learning rates. Even if learning rates can be improved to the Welsh values, the language is still on a downward trajectory; learning rates would need to be significantly higher than in Wales to enable long-term revitalisation of the language in the New Zealand population as a whole.

When the Māori subpopulation is considered separately (Figure 4B), the model predicts that, with current learning rates, the language is on a downward trajectory, but language revitalisation will be successful in the long-term if similar learning rates to those in Wales can be achieved. This is because the initial condition for the Māori subpopulation is more favourable than in the overall New Zealand population (see Table 1). Model predictions for the non-Māori subpopulation (Figure 4C) are similar to those for the population as a whole (Figure 4A) because non-Māori make up the majority of the total population.

The prediction that the language is on a downward trajectory in the Māori subpopulation is consistent with census data showing a decline on the proportion of Māori under the age of 24 who are able to speak the language from 21% in 2001 to 16% in 2013 (18, 61). However, data from the survey of Māori wellbeing show an increase in the proportion of Māori under the age of 24 able to speak more than a few words or phrases from 42% in 2001 to 55% in 2013 (62). These differences could be due to different respondent populations or different categorisations of self-assessed proficiency levels. The survey of Māori wellbeing also showed that the biggest increases in the proportion able to speak the language were in the 25-34 and 35-44 age brackets (62). This suggests that adult learning is a significant pathway to language acquisition and therefore the true learning rate parameters may be higher than the estimates in Table 2, which were derived from statistics on school-level learning.

The calibrated model can be used to evaluate whether the language’s current trajectory will meet stated targets. In 2018, the New Zealand government set two targets: (i) by 2040, 150,000 Māori will speak te reo Māori as a primary language; and (ii) by 2040, one million New Zealanders will be able to speak at least basic te reo Māori (18). Target (i) translates into model variables as 14% of Māori in the proficient category. Achieving target (i) would require increasing the learning rates in the Māori subpopulation to a level approximately 50% higher than the Welsh learning rates. For target (ii), “at least basic” means to being able to speak more than a few words or phrases (18). This does not map as easily onto model variables, as we defined the independent category as being able to speak “fairly well”, which is a higher level of proficiency. If we assume that, of those with “at least basic” competency, half are in the basic category and half are in the independent or proficient categories, target (2) translates to 8.7% of the whole population in the independent or proficient categories. However, even with this relatively generous interpretation, the model predicts that achieving this target would require the learning rates for the population as a whole to be approximately 5 times higher than the Welsh rates. See Supplementary Information for details of target calculations and model predictions.

### Model extension to two interacting population groups

It is likely that reality is somewhere between the scenario shown in Fig. 4A, which assumes a single, homogeneously mixed population, and that in Fig. 4BC, which assumes two completely separate and non-interacting subpopulations. In addition, learning rates in each of the subpopulations are likely to differ significantly, given the influence of cultural exposure and access to language domains on language acquisition and development. Capacity for intergenerational transmission and demographic parameters such as birth and death rates may also differ between ethnicities. The degree of intermixing, as well any variation in transmission and learning rate parameters, will affect model predictions. Addressing this variation would not necessarily lead the model towards the competition approach of (34), but would allow for a more realistic reflection of the development of multicultural and multi-language coexistence.

We propose a more realistic model that splits the total population into two subpopulations, each having three proficiency levels. For simplicity, we assume that both subpopulations are of fixed size and that all individuals belong to the same subpopulation as their parents (i.e. no multi-ethnicity families), but we allow social mixing between subpopulations, controlled by an additional parameter *γ*. In the model, individuals progress to the next proficiency level at a rate that depends on their frequency of interaction with proficient users. If there is no mixing (*γ* = 0), potential learners can only interact with proficient users in their own subpopulation; if there is complete mixing (*γ* = 1), learners interact with all proficient users equally, regardless of which subpopulation they belong to. As with the simple version of the model, interactions with proficient users may span a spectrum from formal teacher-student interactions to language acquisition via informal social contacts. As previously, we do not attempt to model these variations, but simply assume that the more interactions there are between basic/independent users and proficient users, the more learning will take place. The only modification we have made is to control the degree to which interactions take place within subpopulations relative to between subpopulations.

We denote the proportion of the Māori subpopulation in the three proficiency categories as *B*_*M*_, *I*_*M*_, *P*_*M*_ and the proportion of the non-Māori subpopulation as *B*_*X*_, *I*_*X*_, *P*_*X*_. This gives a set of four differential equations:

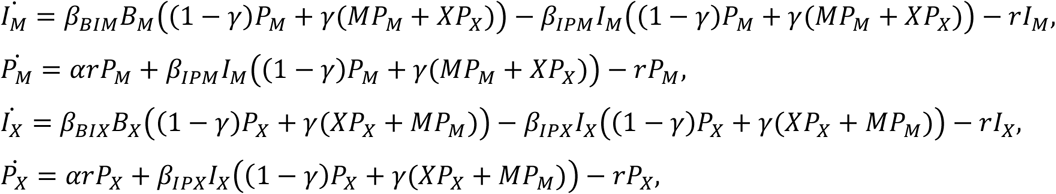

where *M* and *X* are the relative sizes the Māori and non-Māori subpopulations respectively. We assume that *M* = 0.15 and *X* = 0.85 and use the same initial conditions and intergenerational transmission rate as previously. Figure 5 shows the effect of mixing between the two subpopulations, as measured by the parameter *γ*, under the Welsh learning rates. As seen previously in Fig. 4BC, when there is no mixing (*γ* = 0, completely independent subpopulations), the model predicts that te reo Māori will thrive among Māori but die out in the non-Māori subpopulation. However, even a relatively small degree of mixing drastically reduces the predicted long-term number of proficient users among Māori, without producing any significant benefit for non-Māori. This is because the Māori subpopulation has a relatively high proportion of proficient users who promote learning among Māori (either through formal teaching or informal social interactions). However, if these proficient users are spread across the much larger total population, they become too diluted and are unable to sustain the language on an upward trajectory in either subpopulation.

**Figure 5:**
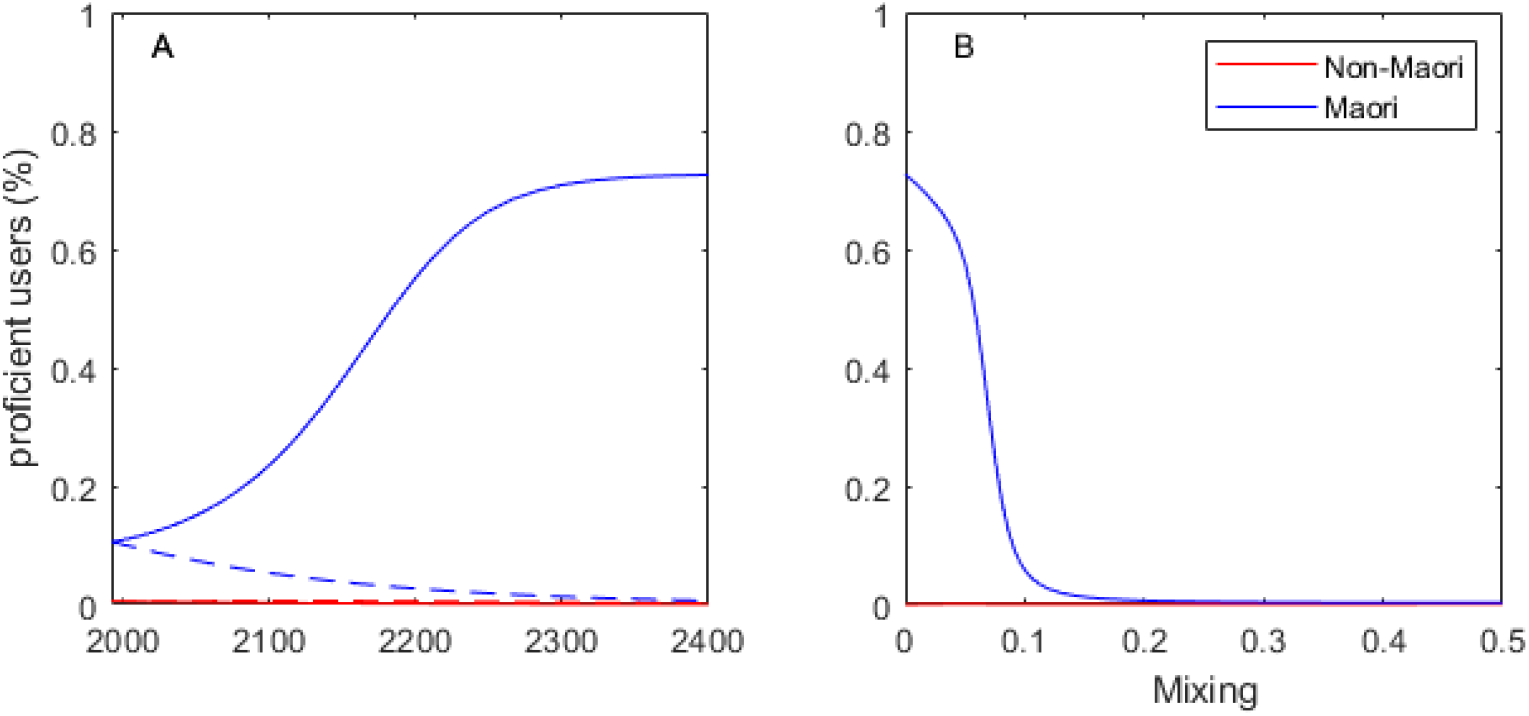
Overuse of Māori teachers in the non-Māori population could have a detrimental effect on long-term language revitalisation. (A) With no mixing of teachers between the populations (*γ* = 0, solid curves) te reo Māori is predicted to thrive in the Māori subpopulation (blue) under Welsh learning rates, but decline to extinction in the non-Māori subpopulation (red). With fully homogeneous mixing of teachers (*γ* = 1.0, dashed curves), te reo Māori is predicted to go extinct in both subpopulations. (B) Even a small degree of mixing of teachers between populations (*γ* > 0) drastically reduces the long-term number of proficient users in the Māori subpopulation (blue), without having any significant beneficial effect for the non-Māori subpopulation (red).

## Discussion

Prevalent themes among language revitalisation strategies include increasing knowledge, skill and proficiency across the population and creating the conditions that support the sustenance of intergenerational transmission (16, 18, 20, 41). The strategies ultimately aspire to develop bilingual or multilingual populations where languages can coexist rather than compete (46, 63). We have developed a dynamic model that considers a population in terms of levels of proficiency with an endangered language. This simple model can offer mathematical validation of language revitalisation strategies and aligns well with their key objectives and cultivation of bilingualism.

The model predicts that there are two possible long-term outcomes for an endangered language, depending on the average rate at which individuals progress from one proficiency level to the next. If the learning rates are too low, the language will be on a trajectory towards extinction; if the learning rates are sufficiently high, the language will be on a trajectory to revitalisation. The threshold value of the learning rates needed to ensure successful revitalisation depends on the number of current language users via the model’s initial condition. We used data from the 2013 New Zealand census to determine these initial conditions. These values are approximate because the data come from a self-assessment of proficiency (18), which can be biased by factors such as social and cultural identity, language anxiety, and the purpose of the survey itself (64, 65). In addition, there were contextual and administrative differences between the 2001 and 2013 surveys (62) and the self-assessment levels do not map perfectly to the proficiency categories in the model. If, as suggested by some studies (65) and consistent with the Māori culture value of whakaiti (humbleness), self-assessed proficiency levels tend to be underestimates, our estimates for the threshold learning rates required for language survival would be conservative. However, better quality data or additional time points, would be needed to confirm this. One advantage of the model is that the qualitative predicted outcome (extinction or revitalisation) is not sensitive to the exact quantitative values of parameters and initial conditions, provided these are not close to the threshold level.

We estimated the learning rate parameters in the model using a combination of data on trends in the number of speakers of Welsh, and the relative rates of participation in language learning at different school levels. With the estimated learning rates for te reo Māori, the model predicts that the language is currently on a downward trajectory within the Māori population and will not meet government targets by 2040 without a major increase in learning rates to levels above those achieved in Wales.

We have used the model to explore language revitalisation strategies aimed at increasing the learning rate parameters of the model. These parameters represent the average rate at which individuals progress from one proficiency level to the next, when taught by a proficient language user. The education system is recognised as one of the most powerful levers available to government policymakers for the acquisition of an endangered language (18). Statistics broken down by age brackets (62) suggest that adult learning of te reo Māori is relatively strong and school-age learning lags behind. This suggests that strategies targeting learning at schools are likely to have the highest potential benefits. Possible government measures include the provision of language-medium early childhood education; integration of the language into the primary and secondary school curriculum; development of the quantity and quality of teachers; investment in language-immersion education as a crucial avenue for the development of proficiency. These have all contributed to the nascent Welsh language revitalisation (24, 55). Other factors in the Welsh case include the availability of university-level and adult education; the struggle to achieve equal recognition and usage of Welsh in the public and institutional spheres; increases in Welsh publishing, broadcast and web media, and software (24).

The model shows that if proficient teachers, who are predominantly Māori, are spread across the whole population, this is detrimental to the language trajectory in the population as a whole, because the limited pool of teachers is spread too thinly. This relates directly to New Zealand government policy, which has a dual strategy of Maihi Māori (22), a by-Māori, for-Māori language revitalisation strategy, and Maihi Karauna (18), the Crown strategy for revitalisation at a national level. Our results suggest that resources should be focused on the Maihi Māori aim of supporting whānau (families) and iwi (tribes) to realise te reo as an everyday language. . This does not mean that learning among non-Māori is unimportant or should not be supported, but that where capacity is limited by the number of teachers, learning among Māori should be prioritised initially. If and when the language is determined to be on a healthy trajectory towards revitalisation among the Māori population, resources and teachers can be distributed more broadly to promote learning across the whole population.

Another potential route to success would be a significant increase in the intergenerational transmission rate, which is the other key parameter of the model. However, in reality, this is likely to be a slowly varying parameter because an increase requires increased intergenerational participation and continuation of language use, and so progress may only be evident over multiple generations (21). Official statistics show that the biggest uptake in the language between 2001 and 2013 was in the 25-44 age bracket (62), which is the age group most likely to be raising young children. Hence, there may be synergistic effects of increasing learning rates and intergenerational transmission, which is consistent with the family-focused approach of (21).

The model has the potential to provide quantitative numerical targets for proficiency levels, which may be used to inform language revitalisation policy and implementation, and evaluate its effectiveness. Furthermore, it is likely that the social norms and values in which such strategies take place are at least as important as the strategies themselves. Attitudes towards te reo Māori among younger generations are shifting, and the government has specifically identified the importance of New Zealanders recognising the value of te reo Māori as a key element of the national identity (18). These factors may also be affected by changing demographics. For example, in New Zealand, the relative size of the Māori population is predicted to grow from 15% to 18% over the next 20 years (60).

The model neglects a range of factors that are likely to be important for the rates of language use and language learning. This is appropriate given the limited data available, and allows the model to focus on comparing alternative strategies or interventions. The model could be extended, for example by making the model spatially explicit to allow for geographic variations in language proficiency levels. This has been done in models of language competition via use of a diffusion term to model localised interactions among individuals (35, 36), which has successfully modelled historical data on language shift (38, 40). The three categories in the model are a coarse classification scale for language proficiency. A finer scale could be used by adding more proficiency levels, though this would ideally require appropriate data to estimate parameter values and initial conditions.

There are potential synergies between the approach we have taken and the literature on cultural evolution, which is the theory that cultural change can be viewed as a Darwinian process involving variation, selection and transmission of cultural traits (32, 54). Reaction-diffusion equations have been used extensively to model the uptake and spatial spread of new technologies and ideas (43). Our model has at its core a social learning mechanism – a key concept in cultural evolution meaning the non-genetic transmission of information from one individual to another (33). Several social learning biases have been documented, including conformity bias, where people are more like to copy popular traits, and prestige bias, where people are more likely to copy traits of successful individuals (54). These biases could play a role in people’s decisions about whether to learn an endangered language. Conformity bias would accelerate learning rates as average proficiency levels increase, which could be modelled via a nonlinearity in the per capita transition rates. Prestige bias would depend on the perceived status (e.g. wealth, political power) of language users, which could be modelled by introducing a status variable (34, 42). These biases and other changes in attitudes towards the language could move the trajectory away from model predictions as it moves outside the conditions in which the calibration data were collected. The model will need refinement and recalibration as these effects play out and there may much to be learned by combining our approach with these and other concepts from cultural evolution.

## Author Contributions

TBW conducted the literature review, designed the methodology, collated census and survey data, carried out numerical solutions of the model, and wrote the first draft of the manuscript. MJP conceived the study, designed the methodology, contributed to the literature review, carried out numerical solutions of the model, produced graphs for the manuscript and critically revised the first draft. RKM and DH contributed to the literature review and to revising draft versions of the manuscript. AJ conceived the study, designed the methodology, carried out numerical solutions of the model, produced graphs for the manuscript and critically revised the first draft. All authors approved the final version of the manuscript.

## Acknowledgements

The whakataukī (Māori proverb) in the title of this paper was adapted from He inoi kia mau te reo Māori (a prayer for the Māori language) by Tā Kīngi Matutaera Īhaka. The authors are grateful to Mary Boyce (Kaihautū Ako Māori, University of Canterbury) and Dr Abby Suszko (Kaiārahi Māori, University of Canterbury) for useful discussions about this research project. MJP and AJ were partly supported by Te Pūnaha Matatini.

## Data Accessibility

This article does not report new data.

## Ethics

This article does not present research with ethical considerations.

