## Supplementary Information for "Kia kaua te reo e rite ki te moa, ka ngaro: Do not let the language suffer the same fate as the moa"

**New Zealand government targets**

We used the model to investigate the learning rate parameters that would be required to meet two targets set by the New Zealand government in 2018 (Te Puni Kōkiri, 2018):

1. By 2040, 150,000 Māori will speak te reo Māori as a primary language.
2. By 2040, one million New Zealanders will be able to speak at least basic te reo Māori.

To translate these targets into model variables, we first converted the target numbers in percentages of the relevant population. We did this using published population projections for 2038 for the Māori and total populations of New Zealand (StatsNZ, 2013c) – see Supplementary Table S1. In addition, for target 2, we assumed that, of the 17.3% of the population able to speak at least basic te reo Māori, half are in the basic category and half are in the independent or proficient category. This leads to a target of 8.7% in the independent or proficient category.

To evaluate target 1, we ran the model using initial conditions for the Māori subpopulation shown in Table 1 of the main text. To evaluate target 2, we ran the model using initial conditions for the overall population shown in Table 1 of the main text. In each case, we varied the learning rates $\beta_{BI}$ and $\beta_{IP}$ (keeping the ratio $\beta_{BI}/\beta_{IP}$ constant) to find the values required to meet the target. Supplementary Figure S1a shows the model solution for the Māori population, with learning rates sufficient to meet target 1. These are approximately 1.5 times the Welsh learning rates. Figure S1b shows the model solution for the population as a whole, with learning rates sufficient to meet target 2. These are approximately 5.4 times the Welsh learning rates.

|  | **Target 1**  **Māori population** | **Target 2**  **Overall population** |
| --- | --- | --- |
| Projected population (2038) | 1059400 | 5769800 |
| Government target (2040) | 150000 (14.2%)  use te reo as primary lang | 1000000 (17.3%)  able to speak at least basic reo |
| Model target | 14.2% proficient | 8.7% independent or proficient |
| Learning rates required | 1.5 x Welsh rates | 5.4 x times Welsh rates |

**Supplementary Table S1.** Mapping of New Zealand government targets to model targets. Rows of the table show: population projections for 2038 for the Māori population (target 1) and total population (target 2) of New Zealand, published by StatsNZ (2013c); government targets to be met by 2040; approximate model targets; learning rates required in the model (relative to the estimated Welsh learning rates) to meet the target by 2040.

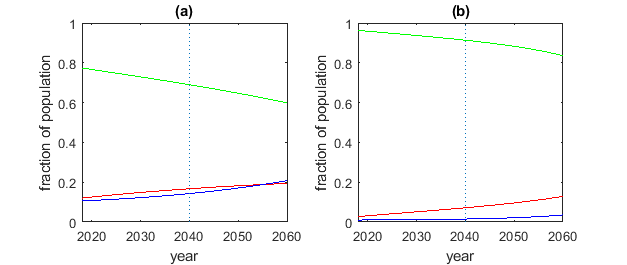

**Supplementary Figure S1.** Model predictions for the proportion of the population in the basic (green), independent (red) and proficient (blue) categories under two scenarios: (a) target 1, the Māori subpopulation with learning rates 1.5 times the Welsh learning rates, leading to 14% in the proficient category in 2040; (b) target 2, the overall population with learning rates 5.4 times the Welsh leading rates, leading to 7.3% in the independent and 1.5% in the proficient category in 2040.
